# Impact of various vaccine boosters on neutralization against Omicron following prime vaccinations with inactivated or adenovirus-vectored vaccine

**DOI:** 10.1101/2022.01.25.476850

**Authors:** Qingrui Huang, Jiawei Zeng, Qingyun Lang, Feng Gao, Dejun Liu, Siyu Tian, Rui Shi, Ling Luo, Hao Wang, Liping Hu, Linrui Jiang, Yawei Liu, Kailiang Li, Yunbo Wu, Junjie Xu, Wenxi Jiang, Ning Guo, Zhihai Chen, Xiaohua Hao, Ronghua Jin, Jinghua Yan, Yufa Sun

## Abstract

Since the first report on November 24, 2021, the Omicron SARS-CoV-2 variant is now overwhelmingly spreading across the world. Two SARS-CoV-2 inactivated vaccines (IAVs), one recombinant protein subunit vaccine (PRV), and one adenovirus-vectored vaccine (AdV) have been widely administrated in many countries including China to pursue herd immunity. Here we investigated cross-neutralizing activities in 341 human serum specimens elicited by full-course vaccinations with IAV, PRV and AdV, and by various vaccine boosters following prime IAV and AdV vaccinations. We found that all types of vaccines induced significantly lower neutralizing antibody titers against the Omicron variant than against the prototype strain. For prime vaccinations with IAV and AdV, heterologous boosters with AdV and PRV, respectively, elevated serum Omicron-neutralizing activities to the highest degrees. In a mouse model, we further demonstrated that among a series of variant-derived RBD-encoding mRNA vaccine boosters, it is only the Omicron booster that significantly enhanced Omicron neutralizing antibody titers compared with the prototype booster following a prime immunization with a prototype S-encoding mRNA vaccine candidate. In summary, our systematical investigations of various vaccine boosters inform potential booster administrations in the future to combat the Omicron variant.

## Introduction

SARS-CoV-2 variant B.1.1.529, first reported on November 24, 2021, is now rapidly spreading across the world, especially in regions where the Delta variant is circulating, suggesting its potential of overtaking Delta to become the next dominant variant. This variant bears up to 37 mutations in the spike protein including 15 within the receptor binding domain (RBD) (*1*) , the primary target of SARS-CoV-2 neutralizing antibodies (*2–4*), which raises immense concern on immune evasion. On November 26, 2021, the World Health Organization (WHO) designated B.1.1.529 as the fifth variant of concern (VOC) and named it Omicron (*5, 6*). Two SARS-CoV-2 inactivated vaccines (IAVs, CoronaVac by Sinovac and BBIBP-CorV by Sinopharm) with a two-dose vaccination regimen, one recombinant protein subunit vaccine (PRV, ZF2001 by Anhui Zhifei Longcom) with a three-dose vaccination regimen, and one single-dose recombinant adenovirus-vectored vaccine (AdV, Convidecia by CanSino) have been given conditional approval for general public use or approved for emergency use by China (*7–11*). These four vaccines form the core of China’s vaccination program. To date, several billion doses of those vaccines have been widely administered in many countries, including China, with the aim of achieving herd immunity against SARS-CoV-2. Moreover, to combat wanning vaccine-elicited immunity with time and emerging variants, several clinical trials with homogenous or heterogenous platform vaccine boosters have also been conducted (*12–14*).

Recent preliminary studies have reported that neutralization elicited by one mRNA vaccine (BNT162b2 by BioNTech in collaboration with Pfizer) is substantially reduced against the Omicron variant (*15, 16*). The Omicron variant also escapes the majority of current therapeutic monoclonal antibodies (*17, 18*). It is urgent to assess residual neutralization levels against the Omicron variant that are afforded by widely used vaccines in China, including IAV, PRV, and AdV; it is equally or even more important to investigate which vaccine booster strategy is able to maximize neutralization capacity against the Omicron variant. In this study, we systematically assessed cross-neutralization activity of human antisera against the Omicron variant elicited by infection or full-course vaccinations with IAV, PRV, or AdV, and by homologous or heterogenous vaccine boosters. All of the currently approved SARS-CoV-2 vaccines use the prototype strain-derived proteins as vaccine immunogens. In a mouse model, we further investigated Omicron-neutralizing activity increase in magnitude induced by various booster vaccine candidates that were developed based on different SARS-CoV-2 variants so as to yield key knowledge about guiding potential booster shots in the future.

## Results

To investigate the Omicron variant’s sensitivity to immunity elicited by infection or full-course vaccination, we measured the binding, blocking and neutralizing activities of serum specimens obtained from 25 convalescent individuals (one month since convalescence), from 30 recipients of two-dose IAV (one month since receipt of the second dose), from 30 recipients of one-dose AdV (one month since receipt of the injection), and from 28 recipients of three-dose PRV (six months since receipt of the third dose) (Table S1). For all of the serum specimens, the binding antibody titers were 5–15 times lower for the Omicron than variant for the prototype strain (Fig. 1A). Notably, except for one specimen from an individual receiving PRV vaccinations scored as positive, all of the other specimens lost the blocking activity against the Omicron variant, whereas nearly all of the specimens exhibited positive blocking activity against the prototype strain (Fig. 1B). Pseudovirus-based neutralization assays demonstrated that the geometric mean NT_50_ titers against the Omicron variant were 20-, 10-, 6-, and 4-fold lower than those against the prototype strain in serum specimens from convalescent patients, IAV recipients, AdV recipients, and PRV recipients, respectively (Fig. 1C). In addition, despite being obtained six months following full-course vaccination, the serum samples from individuals who had received PRV vaccinations had the highest neutralizing NT_50_ titers against the Omicron variant, but these titers were approximately 1/5 of the titer of convalescent sera against the prototype strain (Fig. 1C). In contrast, the neutralizing activity against the Omicron variant afforded by full-course vaccination with IAV or AdV was less than 1/20 of the activity of convalescent sera against the prototype strain (Fig. 1C), suggesting a potential of neutralizing insufficiency against Omicron infection.

**Fig. 1.**
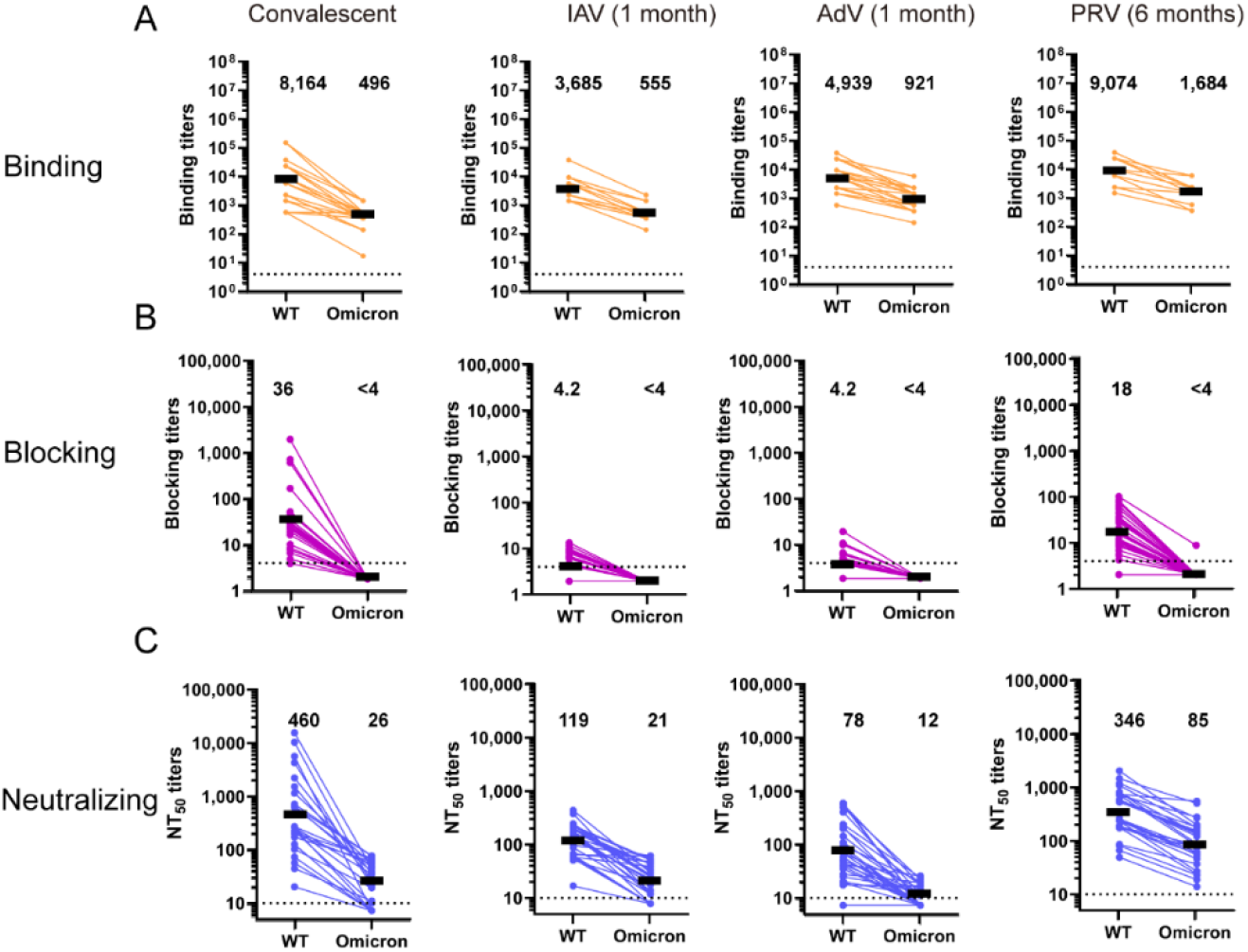
Serum binding, blocking and neutralizing antibody titers of convalescents and vaccinated individuals were markedly lower against Omicron compared to wild-type SARS-CoV-2. Samples included sera obtained from convalescents (n=25), one month after two-dose vaccination with IAV (n=30) or a single-dose AdV (n=30), and six months after three-dose vaccination with PRV (n=28). (A) Serum binding antibody titers were detected by ELISA assay. Coated antigen was wild type RBD or Omicron RBD. Dotted line indicates the limit of detection (>8). (B) hACE2-blocking antibody titers were detected by ELISA using wild-type or Omicron spike protein, which binds to human ACE2. The dotted line indicates the limit of detection (>4). (C) Pseudovirus neutralization titers, expressed as 50% neutralization dilution (NT_50_). The pseudoviruses used in the study included both wild-type strain and Omicron. The dotted line indicates the limit of detection (>10). IAV represents inactivated vaccines (CoronaVac and BBIBP-CorV). PRV represents recombinant protein subunit vaccine (ZF2001). AdV represents adenovirus-vectored vaccine (Convidecia).

To explore the impact of a homologous or heterologous booster at 4–8 months following IAV full-course vaccination on vaccine-induced antibodies against the Omicron variant, we obtained 42 serum specimens from participants receiving no vaccine booster, 39 serum specimens from participants receiving homologous IAV booster (IAV-b) and 45 serum specimens from participants receiving heterologous PRV booster (PRV-b) (Table S1). The samples were from a single-center, open-label, randomized controlled clinical trial at Beijing Ditan Hospital. An additional seven specimens from individuals who had received heterologous AdV vaccine booster following two doses of IAVs 4–8 months earlier from Peking Union Medical University Hospital were also included (Table S1). Binding, blocking and neutralizing antibodies titers in all of the groups were markedly higher against the prototype strain than against the Omicron variant (Fig. 2). The booster dose with IAV, PRV, and AdV vaccine induced 3-, 7-, and 40-fold increase in Omicron-binding antibody titers compared with the no-booster control (Fig. 2A). At 4–8 months after full-course vaccination with IAV, Omicron-blocking antibodies were close to or below the lower limit of detection (4-fold dilution of plasma), and the blocking positive rate was only 2% (Fig. 2B and Fig. S1). The IAV, PRV, and AdV vaccine boosters led to an increase of blocking positive rates to 54%, 71%, and 57%, respectively (Fig. 2B and Fig. S1). For neutralizing antibodies against the Omicron variant, the genomic mean NT_50_ titers were below the lower limit of detection (10-fold dilution of plasma) in the control group, whereas the titers rose to 113, 207, and 709 in the IAV, PRV and AdV booster groups, respectively (Fig. 2C and Fig. S2), indicating that the heterologous vaccine booster with AdV was superior to the homologous IAV vaccine booster in improving the neutralizing activity against the Omicron variant.

**Fig. 2.**
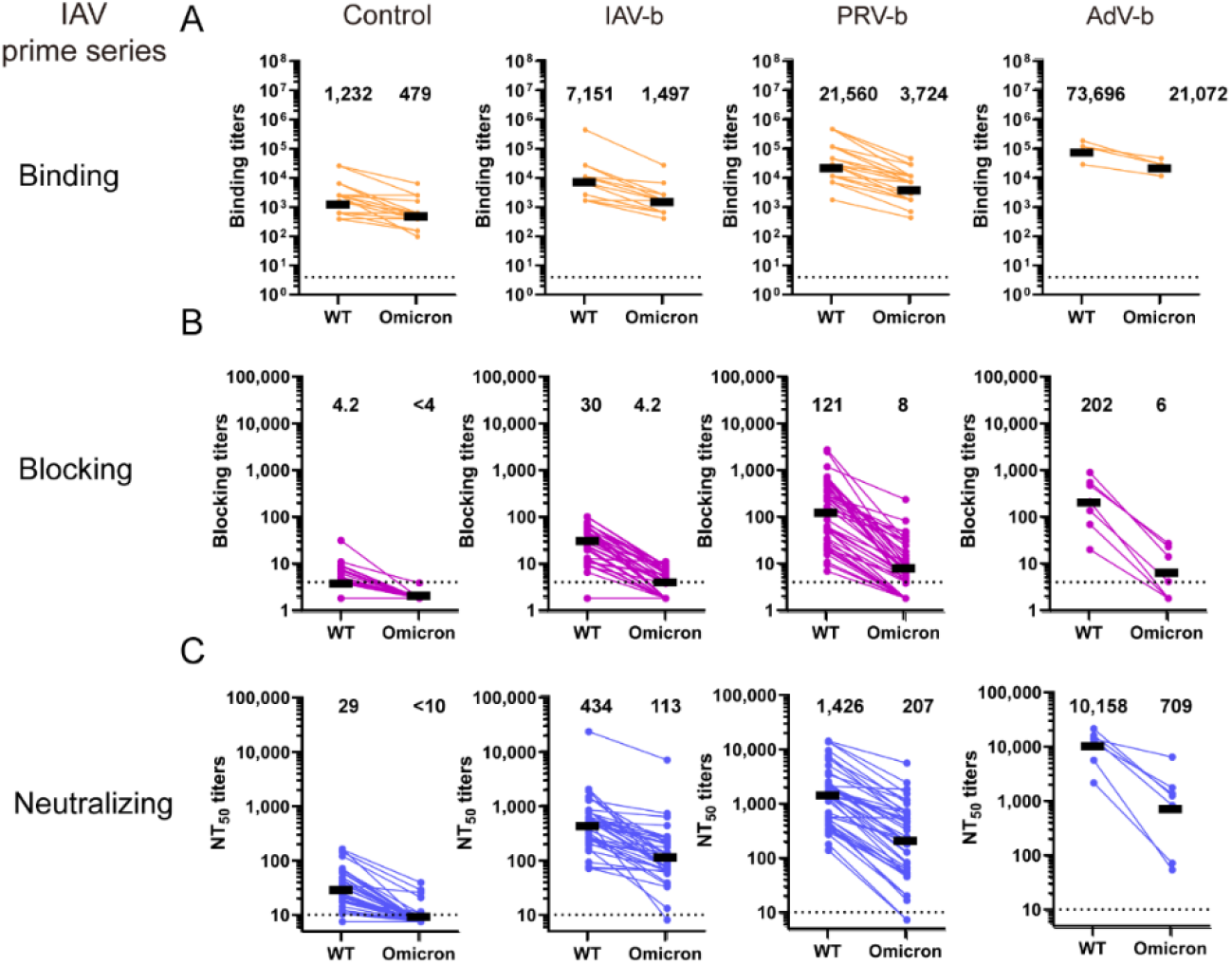
Serum binding, blocking and neutralizing antibody titers of no booster control and various vaccine boosters following prime vaccination with two-dose IAVs. Samples were obtained from participants without vaccine booster (n=42), with IAV (IAV-b, n=39), AdV (AdV-b, n=7), or PRV (PRV-b, n=45) boosters following two-dose prime vaccination with IAV 4-8 months earlier. (A) Serum binding antibody titers were detected by ELISA. Coated antigen was wild type RBD or Omicron RBD. Dotted line indicates the limit of detection (>8). (B) hACE2-blocking antibody titers were detected by ELISA using SARS-CoV-2 wild type and Omicron spike proteins which bind to human ACE2. The dotted line indicates the limit of detection (>4). (C) Pseudovirus neutralization titers, expressed as 50% neutralization dilutions (NT_50_). The pseudoviruses used in the study included wild-type strain and Omicron. The dotted line indicates the limit of detection (>10). IAV represents inactivated vaccines (CoronaVac and BBIBP-CorV). PRV represents recombinant protein subunit vaccine (ZF2001). AdV represents adenovirus-vectored vaccine (Convidecia).

To examine the effect of various booster vaccinations following a single-dose prime vaccination with AdV, we conducted similar tests with serum specimens from individuals receiving no booster injection (control, n=30), IAV booster (n=30), PRV booster (n=30), or AdV booster (n=30) at 4–8 months following the primary AdV vaccination (Table S1). All of the samples were collected one month following booster vaccine injection at Chinese PLA General Hospital. The Omicron-binding antibody titers were boosted 4-, 25-and 16-fold by the booster injection of heterologous IAV and PRV vaccines or homologous AdV vaccine, respectively, compared with the no-booster control (Fig. 3A). For Omicron-blocking antibodies, the PRV and AdV booster groups exhibited an identical blocking positive rate (80%), which was higher than that of the control (3%) or IAV booster group (53%) (Fig. 3B and Fig. S3). The neutralizing NT_50_ titers for the Omicron variant in the control group and the IAV, PRV, and AdV booster groups were 15, 68, 313, and 228, respectively (Fig. 3C and Fig. S4). Notably, a heterologous PRV booster induced the highest degree of neutralizing immunity against both the prototype and the Omicron strains compared with the no-booster control and the other two boosters (Fig. 3C and Fig. S4).

**Fig. 3.**
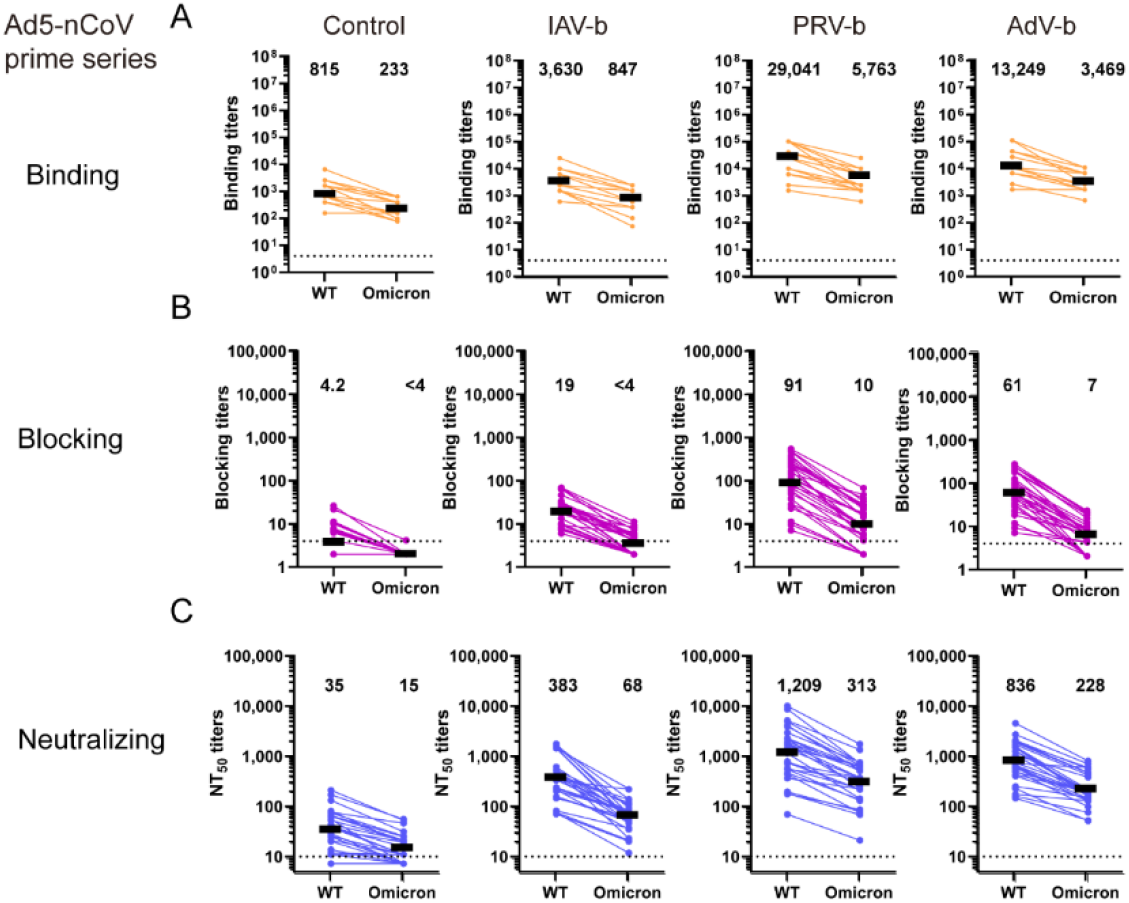
Serum binding, blocking and neutralizing antibody titers of no booster control and various vaccine boosters following a single-dose prime vaccination with AdV. Samples were obtained from participants without vaccine booster (n=30), with IAV (IAV-b, n=30), AdV (AdV-b, n=30), or PRV (PRV-b, n=30) boosters following a single-dose prime vaccination with IAV 4–8 months earlier. (A) Serum binding antibody titers were detected by ELISA. Coated antigen was wild type RBD or Omicron RBD. The dotted line indicates the limit of detection (>8). (B) hACE2-blocking antibody titers were detected by ELISA using SARS-CoV-2 wild type and Omicron spike proteins which bind to human ACE2. The dotted line indicates the limit of detection (>4). (C) Pseudovirus neutralization titers, expressed as 50% neutralization dilutions (NT_50_). The pseudoviruses used in the study included wild-type strain and Omicron. The dotted line indicates the limit of detection (>10). IAV represents inactivated vaccines (CoronaVac and BBIBP-CorV). PRV represents recombinant protein subunit vaccine (ZF2001). AdV represents adenovirus-vectored vaccine (Convidecia).

All four of the above-mentioned vaccines have been developed based on the prototype strain. It is possible to achieve a superior Omicron-neutralizing immunity by variant vaccine boosters, especially when taking into account that some variants such as Beta and Delta share some common key mutations with the Omicron variant. Next, we developed various RBD-encoding mRNA vaccine candidates based on prototype, Beta, Delta and Omicron strains as booster shots in BALB/c mice following a single-dose prime injection with a prototype S-encoding mRNA vaccine candidate, in order to assess the impact on humoral immunity against the Omicron variant (Fig. 4A). Beta-Delta represents a vaccine combination of Beta and Delta vaccine candidate with either half the other vaccine booster dose. The mice immunized with a prime injection induced binding and neutralizing but no detectable blocking antibodies against the Omicron variant, whereas all binding, neutralizing and blocking antibodies against the prototype were detected in those mice (Fig. 4B-D). The mice immunized with all types of booster shots had significantly elevated anti-prototype strain binding, blocking, and neutralizing antibody titers, as well as anti-Omicron binding and neutralizing antibody titers (Fig. 4B-D). In addition, all of the booster groups induced significantly higher anti-Beta and anti-Delta binding antibody titers compared with the no-booster control group (Fig. S5-6). However, only mice immunized with Delta and Omicron boosters developed significantly higher anti-Omicron blocking antibody titers compared with no booster control (Fig. 4C). Notably, the prototype, Beta, Delta, Beta–Delta and Omicron vaccine boosters elicited Omicron-neutralizing NT_50_ titers with values of 423, 1,202, 3,073, 1,548, and 7,710, respectively, and only those elicited by Omicron booster were significantly higher than those by the prototype booster (Fig. 4D), suggesting that Omicron-based mRNA vaccine booster is superior to prototype-based mRNA vaccine booster in elevating Omicron-neutralizing immunity.

**Fig. 4.**
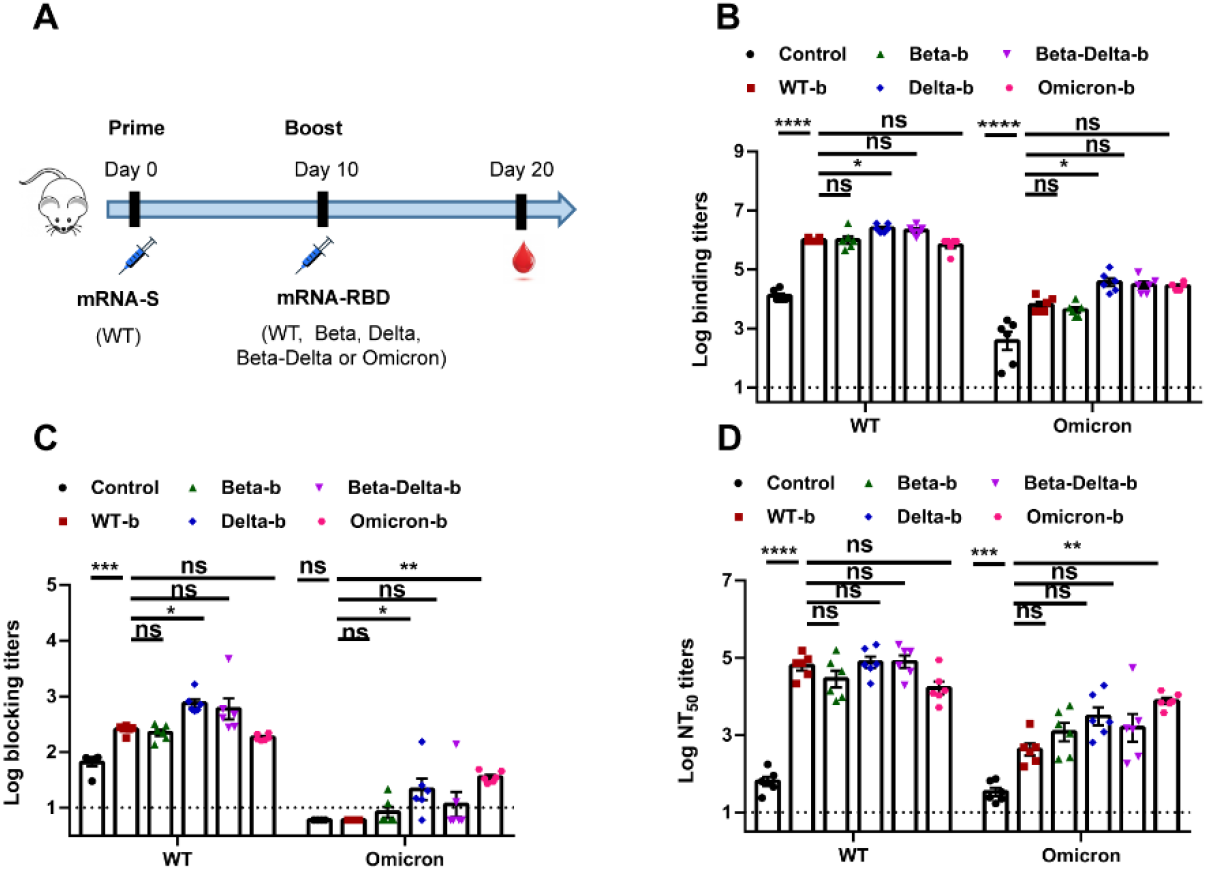
Investigation of various RBD-encoded mRNA vaccine boosters based on SARS-CoV-2 prototype, Beta, Delta, Beta plus Delta and Omicron following a single prime injection. Groups of BALB/c mice (n=6) received no booster or different vaccine booster shot following a prime immunization with wild type S-encoding mRNA vaccine candidate via intramuscular route. Mouse serum were obtained at 10 days following the booster injection. (A) Mice immunization schedule. (B) Mouse sera binding antibody titers were detected by ELISA. Coated antigen was wild type RBD or Omicron RBD. The dotted line indicates the limit of detection (>10). (C) hACE2-blocking antibody titers were detected by ELISA using SARS-CoV-2 wild type and Omicron spike proteins which bind to human ACE2. The dotted line indicates the limit of detection (>10). (D) Pseudovirus neutralization titers, expressed as 50% neutralization dilutions (NT_50_). The pseudoviruses used in the study included wild-type strain and Omicron. The dotted line indicates the limit of detection (>10). P values were analyzed with One-Way ANOVA (ns, p>0.05, *p<0.05, **p<0.01, ***p<0.001, and ****p<0.0001).

## Discussion

Since its emergence, the SARS-CoV-2 Omicron variant has been spreading at an unprecedented speed. Consistent with its far more mutations in spike protein than other variants, we demonstrated that the Omicron variant remarkedly escaped from neutralizing antibody response elicited by full-course vaccinations of approved IAV and AdV vaccines. To boost anti-Omicron response to sufficiently high titers so as to provide some protection against Omicron infection, a booster shot may be necessary. Here we showed that for prime vaccinations with IAV and AdV, a heterologous vaccine booster with AdV and PRV, respectively, generated the highest increase in Omicron-neutralizing antibody titers. Although the sample number in AdV booster with prime IAV vaccinations was small (n=7), the robust increase in the Omicron-neutralizing activity supports the heterologous AdV booster administration. Currently, all of the approved vaccines have been developed based on the SARS-CoV-2 prototype strain. Using the mouse model, we further demonstrated that prototype booster significantly increased the prototype vaccine-elicited Omicron-neutralizing activity. Although the Beta and Delta variants harbor some identical or similar key mutations to the Omicron variant, it is only the Omicron-based but not Beta- or Delta- based vaccine booster that exhibited a significantly higher ability of improving Omicron-neutralizing immunity. Taken together, our systematical investigation of impact of various vaccine boosters on improving Omicron-neutralizing immune response yielded important data to guide possible future booster shots to contain the Omicron pandemic.

Structural comparisons have allowed us to classify RBD-targeted neutralizing antibodies into four class groups (*19*). Class 1 and Class 2 antibodies bind on, or in close proximity to the ACE2-binding footprint, and they can potently neutralize viruses by blocking the interaction of RBD with ACE2, thereby preventing viral attachment to host cells (*20*). As there are concentrated mutations on the ACE2-binding footprint in Omicron RBD, these antibodies showed dramatic or complete loss of the neutralizing activity against the Omicron variant (*17*). Thus, Omicron-blocking titers of human serum specimens showed a substantial decrease compared with the prototype-blocking titers. There are also class 3 and class 4 neutralizing antibodies which bind distant from ACE2-binding site and do not block ACE2 interaction. These antibodies may destabilize the spike trimmer protein, and the antibody epitopes are more conserved in the Omicron variant (*17*). In theory, these antibodies are the main contributors of cross-neutralization against the Omicron variant in prototype-vaccinated human serum. That also leads to a moderate decrease in Omicron-neutralizing antibody titers in comparison with Omicron-blocking antibody titers.

## Acknowledgments

We thank professor Weijin Huang from National Institutes for Food and Drug Control (NIFDC) for kindly providing SARS-CoV-2 prototype and Omicron pseudovirus.

## Funding

This work was supported by the Strategic Priority Research Program of the Chinese Academy of Sciences (XDB29040201) and the National Natural Science Foundation of China (NSFC) (81901680).

## Author contributions

J.Y., Y.S., and Q.H. designed the study; F.G., D.L., Y.L., K.L, Y.W., J.X., W.J., X.H., Z.C., and R.J. collected human serum specimens; J.Z, Q.L., S.T., L.L., H.W., L.H., and L.J. conducted all assays. J.Y., Q.H., and N.G. analyzed and interpreted the data. Q.H. wrote the manuscript. Q.H. and J.Y. discussed and edited manuscript.

## Competing interests

The Institute of Microbiology, Chinese Academy of Sciences (IMCAS) holds the patent on ZF2001 vaccine.

## Data and materials availability

All data are available in the main text or the supplementary materials.

## Materials and Methods

### Human serum samples

Human serum specimens were collected at Peking Union Medical College Hospital, Beijing Ditan Hospital, 309 Hospital of the Chinese People’s Liberation Army, Fangzhuang Community Health Service Center, and the Institute of Microbiology of the Chinese Academy of Sciences (IMCAS). The samples were selected based on availability, and the specimens were obtained from individuals of different genders with no specific inclusion/exclusion criteria. Participants included convalescents from COVID-19, participants who had received full-course vaccination with inactivated vaccines, recombinant protein subunit vaccines, and adenovirus-vectored vaccines, as well as various types of vaccine boosters.

### Ethics statement

This study was reviewed and approved by IMCAS (APIMCAS2021159). This study was conducted in strict accordance with the recommendations in the “Guide for the Care and Use of Laboratory Animals” issued by the Ethics Committee of IMCAS. Informed consent was obtained from all of the participants.

### Cells, pseudoviruses, and animals

HEK293T (ATCC, CRL-1573) cells and Vero E6 cells (ATCC CRL-1586) were cultured at 37°C in Dulbecco’s modified Eagle’s medium (DMEM) supplemented with 10% fetal bovine serum (FBS). Pseudovirus of the SARS-CoV-2 wild type strain and the Omicron variant were provided by Professor Weijin Huang from National Institutes for Food and Drug Control. Specific-pathogen-free (SPF) BALB/c 6-8-week-old female mice were purchased from Beijing Vital River Animal Technology

Co., Ltd (licensed by Charles River). All of the mice used in this study are were housed and bred in a temperature-, humidity- and light cycle-controlled SPF mouse facilities in IMCAS (20±2°C; 50±10%; light, 7:00-19:00; dark, 19:00-7:00).

### Protein expression and purification

The objective recombinant protein sequences carried on the pCAGGS vector were expressed via HEK293T cells. The supernatant of cell culture was collected five days post-transfection. Initially, a Histrap excel 5 mL column (GE Healthcare) was used to isolate proteins. The medium of samples collected from the Histrap columns was substituted with PBS solution and then further purified with the Superdex 200 column (GE Healthcare). Lastly, SDS-PAGE was performed to assess the purity of the protein.

### ELISA

SARS-CoV-2 RBD monomer proteins were coated with 50 mM carbonate-bicarbonate buffer (pH 9.6) in Corning^®^ 96-well Clear Polystyrene Microplates at 200 ng per well. The microplates were blocked by 5% skimmed milk at 37°C for one hour, and then the milk was discarded. The microplates were incubated with 100 μL two-fold serially diluted mice serum at 37°C for one hour. The HRP-labeled anti-mouse Fc secondary antibody (Yeasen) was added after washing the microplates three times. Then, 50 μL of 3, 3′, 5, 5′-tetramethylbenzidine (Beyotime Biotechnology) was used as a substrate and 50 μL of 2 M sulphuric acid was used to stop the reactions. The absorbance was measured at 450 nm using a microplate reader (PerkinElmer). The end-point antibody titers were defined as the highest dilution of the serum that produced an optical absorption value (OD_450_) 2.1 times higher than the background value.

### hACE2-receptor-blocking assay

hACE2-receptor-blocking antibodies were determined by ELISA. Corning® 96-well Clear Polystyrene Microplates were coated with 20 μg/mL human ACE2 (hACE2) protein overnight at 4°C. Serially diluted sera from groups of immunized mice or humans was added into coated wells and then 50 ng/mL of histidine-tagged SARS-CoV-2 S proteins was added into wells for two hours at 37°C. Meanwhile, a set of negative control without S protein and a set of positive control without serum were necessary. After incubation and washing five times, Anti-His-tag-HRP was added and incubated for one hour. Then, 50 μL of 3, 3′, 5, 5′-tetramethylbenzidine (Beyotime Biotechnology) was used as a substrate and 50μL of 2M sulphuric acid was used to stop the reactions. The absorbance was measured at 450 nm using a microplate reader (Perkin Elmer). The reciprocal of the highest serum dilution that resulted in 50% inhibition of receptor binding was used as the titer of the serum.

### Pseudovirus neutralization assay

96 Well White Plates (WHB) were used to detect the neutralization potency. Sera from groups of mice and humans were serially diluted in DMEM medium supplemented with 10% FBS. Pseudoviruses were diluted to 2 × 10^4^ TCID_50_/mL using DMEM (10% FBS), and then 50 μL of the diluted pseudoviruses was added to each well. Meanwhile, a set of negative controls with only medium and a set of positive controls with only pseudovirus were necessary. After incubation at 37°C for one hour, 2 × 10^4^ Vero E6 per cell were added to each well to make a final volume of 200 μl, which was incubated for 24 hours in a 37°C, 5% CO2 incubator. After incubation, the supernatant in the wells was discarded and 100 μL of luciferase detection reagent (PerkinElmer, Inc.) was added. The reaction was shaken at room temperature for two minutes and the fluorescence values were read in a chemiluminescence detector (Promega GloMax). The neutralization NT_50_ titer was defined as the fold-dilution of serum necessary for 50% inhibition of luciferase activity in comparison with virus control samples.

### mRNA production

mRNA was produced using T7 RNA polymerase on linearized plasmids (synthesized by Genescript) encoding codon-optimized SARS-CoV-2 S6P protein or prototype, Beta, Delta, or Omicron RBD glycoprotein (residues 319-541). The mRNA was transcribed to contain a 104 nucleotide-long poly(A) tail, and 1-methylpseudourine-5’-triphosphate was used instead of UTP to generate modified nucleoside-containing mRNA. The mRNA was purified by overnight LiCl precipitation at −20°C, centrifuged at 18,800×g for 30 min at 4°C to pellet, washed with 75% EtOH, centrifuged at 18,800×g for 1 min at 4°C, and resuspended in RNase-free water. The purified mRNA was analyzed by agarose gel electrophoresis and stored at −80°C until use.

### Lipid-nanoparticle encapsulation of mRNA

mRNA was encapsulated in LNPs using a self-assembly process in which an aqueous solution of mRNA at pH=4.0 was rapidly mixed with a solution of lipids dissolved in ethanol. LNPs used in this study contained an ionizable cationic lipid, phosphatidylcholine, cholesterol, and PEG-lipid at a ratio of 50:10:38.5:1.5 mol/mol and were encapsulated at an mRNA to lipid ratio of around 0.05 (wt/wt). The formulations were then diafiltrated against 100 x volume of Phosphate Buffered Saline (PBS) through a tangential-flow filtration (TFF) membrane with 10 kD molecular weight cut-offs (Sartorius Stedim Biotech), concentrated to a required concentration, passed through a 0.22μm filter, and stored at 4°C with a concentration of RNA of about 1 mg/mL.

### Animal experiments

For immunization, 6-8-week-old female BALB/c mice were primarily vaccinated with 4 μg S-encoding mRNA vaccine candidate via intramuscular (i.m.) route. At 10 days following primary immunization, booster injections with PBS as negative control or different variant RBD-encoding mRNA vaccine candidates were administered with a dose of 4 μg. Serum samples were collected at 10 days following booster vaccination, inactivated at 56°C for 30 min, and stored at −80°C until use.

### Statistical analysis

All of the data are expressed as the mean ± standard error of the mean. For all of the analyses, P values were obtained from Student’s t-test (unpaired, two tailed) or One-way ANOVA test. All of the graphs were generated with GraphPad Prism version 7.0 software.

---

**Fig. S1.**
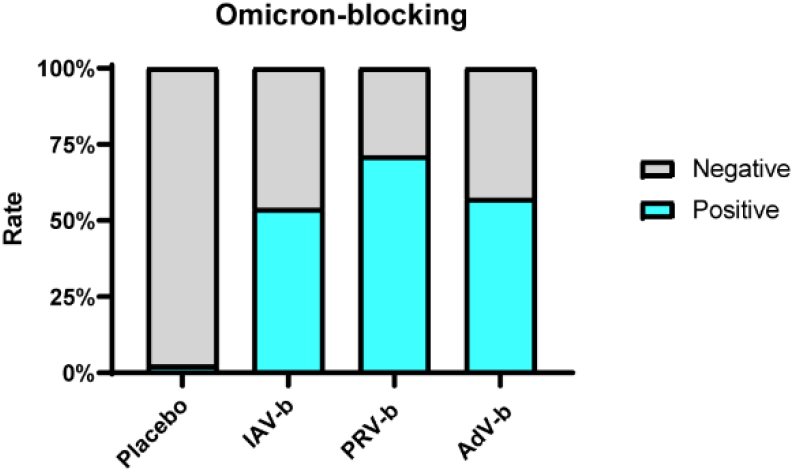
Rate of positive samples for hACE2 blocking antibody detection against Omicron, relative to Fig. 2B. Samples were obtained from participants without vaccine booster (n=42), with IAV (IAV-b, n=39), AdV (AdV-b, n=7), or PRV (PRV-b, n=45) boosters following two-dose prime vaccination with IAV 4–8 months earlier.

**Fig. S2.**
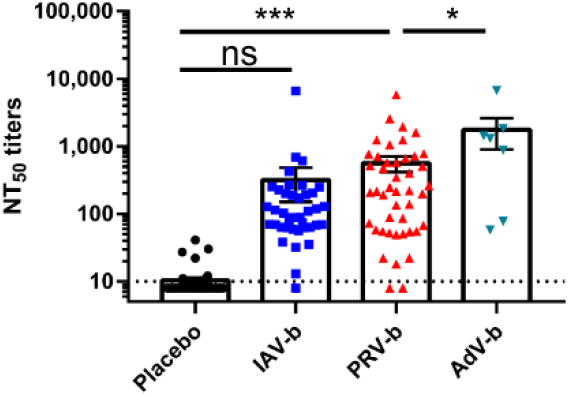
Neutralization titers against Omicron pseudovirus of non-boosted and boosted with different vaccines following previously vaccinated with IAVs, relative to Fig. 2C. Samples were obtained from participants without vaccine booster (n=42), with IAV (IAV-b, n=39), AdV (AdV-b, n=7), or PRV (PRV-b, n=45) boosters following two-dose prime vaccination with IAV 4–8 months earlier. The values were representative of mean ± SEM. P values were analyzed with Student’s t-test (ns, p>0.05, *p<0.05, **p<0.01, ***p<0.001, and ****p<0.0001).

**Fig. S3.**
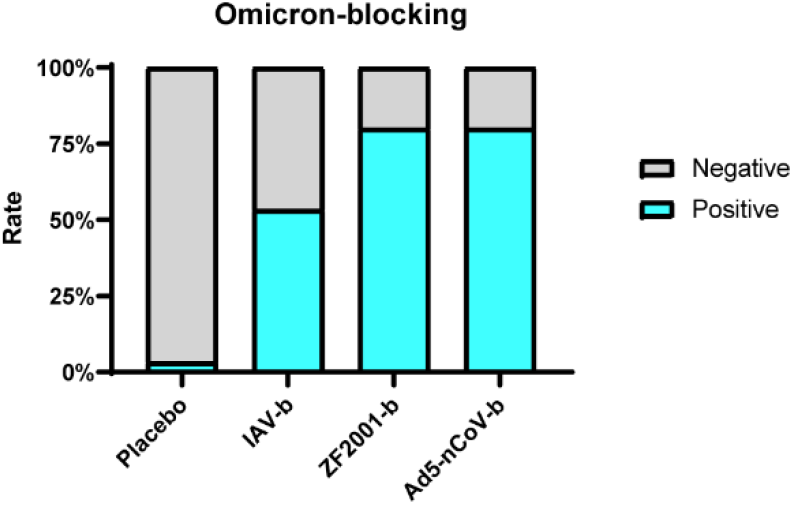
Rate of positive samples for hACE2 blocking antibody detection against Omicron, relative to Fig. 3B. Samples were obtained from participants without vaccine booster (n=30), with IAV (IAV-b, n=30), AdV (AdV-b, n=30), or PRV (PRV-b, n=30) boosters following a single-dose prime vaccination with IAV 4–8 months earlier.

**Fig. S4.**
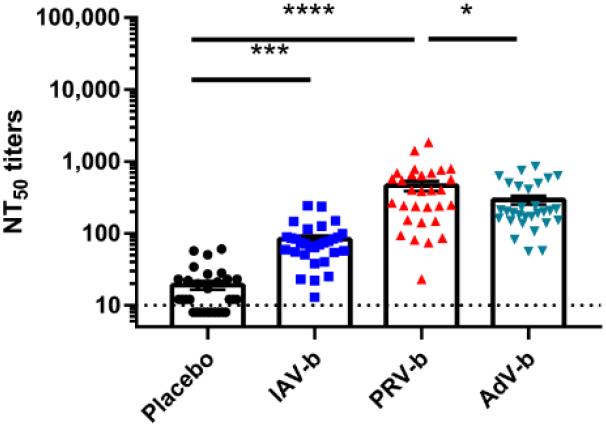
Neutralization titers against Omicron pseudovirus of non-boosted and boosted with different vaccines following previous vaccination with AdV, relative to Fig. 3C. Samples were obtained from participants without vaccine booster (n=30), with IAV (IAV-b, n=30), AdV (AdV-b, n=30), or PRV (PRV-b, n=30) boosters following a single-dose prime vaccination with IAV 4–8 months earlier. The values represent mean ± SEM. P values were analyzed with t-test (ns, p>0.05, *p<0.05, **p<0.01, ***p<0.001, and ****p<0.0001).

**Fig. S5.**
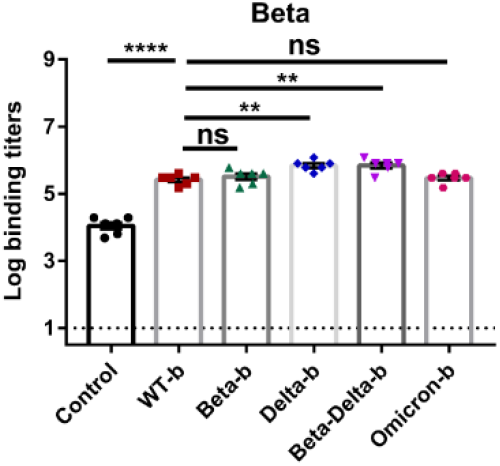
Beta-binding titers of various RBD-encoded mRNA vaccine boosters based on SARS-CoV-2 prototype, Beta, Delta, Beta plus Delta and Omicron following a single prime injection, relative to Fig. 4. The dotted line indicates the limit of detection (>10). P values were analyzed with One-Way ANOVA (ns, p>0.05, *p<0.05, **p<0.01, ***p<0.001, and ****p<0.0001).

**Fig. S6.**
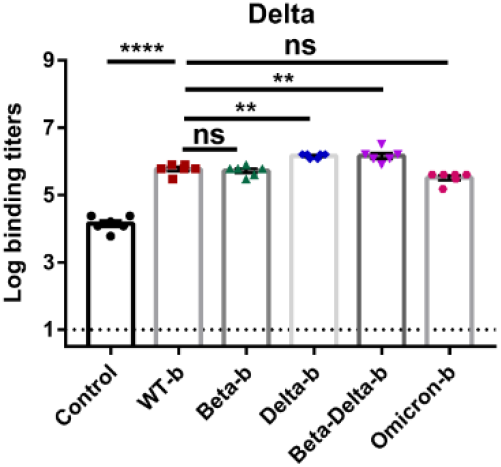
Delta-binding titers of various RBD-encoded mRNA vaccine boosters based on SARS-CoV-2 prototype, Beta, Delta, Beta plus Delta and Omicron following a single prime injection, relative to Fig. 4. The dotted line indicates the limit of detection (>10). P values were analyzed with One-Way ANOVA (ns, p>0.05, *p<0.05, **p<0.01, ***p<0.001, and ****p<0.0001).

**Table S1:**
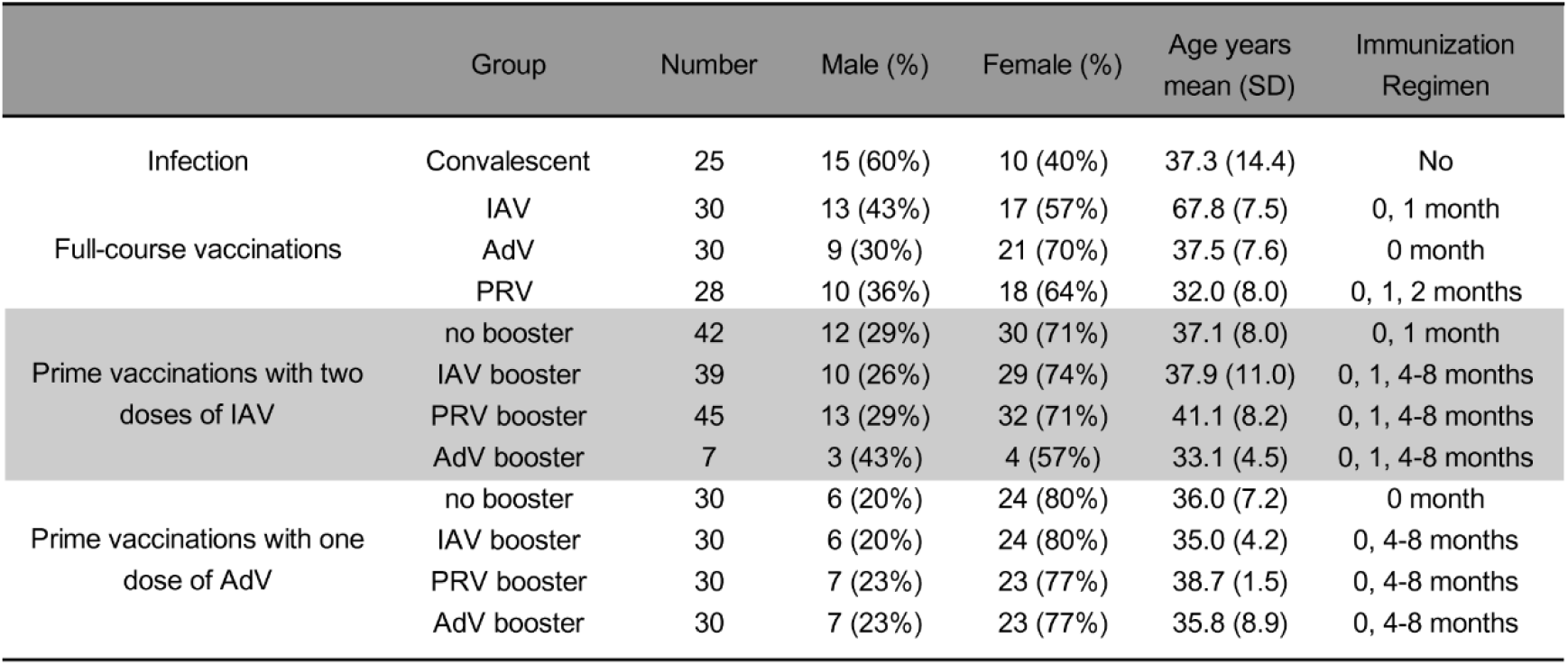
Characteristics of participants

